# STEREOLOGICAL EVALUATION OF TISSUE PRESERVATION AFTER NEUROPROTECTIVE TREATMENTS FOR TRAUMATIC SPINAL CORD INJURY

**DOI:** 10.1101/2022.05.05.490720

**Authors:** David Reigada, Vanesa Soto, María González-Rodríguez, María Asunción Barreda-Manso, Altea Soto, Teresa Muñoz-Galdeano, Rodrigo M. Maza, Manuel Nieto-Díaz

## Abstract

Spinal cord injury (SCI) is a major cause of permanent disability and its causes and pathophysiological effects are very variables between patients. The assessment of tissue damage extent and neurodegeneration degree correlated with the functional evaluation are the most accepted tools to diagnose and prognose the trauma severity. Animal models of SCI have been used for treatment development and in the present work we evaluate the potency of stereological tools to estimate damage degree for a diagnostic of neural degeneration and locomotor and sensorial disability after SCI and the efficacy of different types of therapeutic strategies.

## INTRODUCTION

Two to 7 million people around the world live with permanent disabilities caused by spinal cord injury (SCI) [1,2]. The personal, medical, and social costs of SCI are enormous, including lifetime costs per patient estimated in $1.2 to 5 million for USA [3].

Trauma to the spinal cord exposes neural tissue to forces that depending on its severity, leads to damage in the cell integrity and death among neural and non-neural components of the spinal cord. This primary mechanical damage triggers a stereotyped sequence of secondary events that affect the nervous, blood, and immune systems and involve a wide range of processes among which cell death is prominent (for a comprehensive review of spinal cord pathophysiology see [4]).During the secondary damage, spinal cells surviving the initial mechanical insult become exposed to a deleterious environment that extends neurodegeneration throughout the central nervous system (CNS) [5].

Neurodegeneration largely determine loss of sensory-motor and autonomic functions below the injury level but also affects supraspinal networks leading to additional pathological outcomes such as neuropathic pain [4-6]. The central role that neurodegeneration plays in SCI pathophysiology and in the resulting disabilities identifies it as a major therapeutic target for SCI therapies [7]. It also makes its quantification the gold-standard for diagnosis and prognosis together with functional evaluation through the ASIA exam [4,8].

Whereas magnetic resonance imaging (MRI) is the gold standard to measure neurodegeneration in the clinical practice due to its high correlation found with pathophysiological and functional deficits and severity for spinal cord injuries [9-11], histological techniques are still prominent in animal models of SCI due to the availability of tissue, its accessibility compared to RMI, and the broad range of analyses it may support (anatomical, cellular, RNA, proteins, …) to characterize neurodegeneration.

Staining of myelinated tissue in spinal cord sections by eriochrome cyanine (EC) followed by the Cavalieri’s stereological estimation for white matter preservation is an easy and broadly employed technique to measure neurodegeneration. In previous works we have employed this methodology to estimate tissue preservation in different animal models, SCI severities and neuroprotective treatments [12-14] under the assumption that the resulting estimation of the tissular damage highly correlates with the physiological deficits induced by SCI and the efficacy of any therapeutical strategy.

In the present work, we use the information from these studies and unpublished data to test this assumption and to evaluate the usefulness and limitations of EC-based stereological estimation of tissue damage extent as a reliable biomarker for diagnosis and prognosis in experimental SCI.

## MATERIAL AND METHODS

This article employs data from published and ongoing studies of the Molecular Neuroprotection Laboratory (Hospital Nacional de Parapléjicos). This section provides information on the methods and materials employed in these studies. Further information is available in our previous works [13-15].

### SCI procedure

Animal experimental procedures from the studies described here were in accordance with the European Communities Council Directive (2010/63/EU) and were approved by the Hospital Nacional de Parapléjicos Animal Care and Use Committee (REGA# ES451680000068).

Adult male and female Wistar rats and C57BL/6J mice were employed through these studies. Animals were housed in plastic cages in a temperature and humidity controlled room maintained on a 12:12h reverse light/dark cycle with free access to food and water. Precise information on the experimental conditions can be reached at the original articles [13-15].

For rat models, 12-14 weeks old animals (200 grams) were anesthetized by intraperitoneal injection of sodium pentobarbital (40mg/Kg; Dolethal; Vetoquinol) and xylazine (9 mg/Kg; Xilagesic, Farma Veterinaria) and by subcutaneous injection of the analgesic buprenorfine (0.03 mg/Kg Buprex; Reckitt Benckiser Pharmaceuticals Limited). SCI surgery followed the methodology described in Yunta et al. [15]. Following thoracic vertebra 8 (T8) laminectomy, rats were injured by a mild (150 KDyne) or moderate (200 KDyne) contusion (IH Spinal Cord Impactor, Precision System & Instrumentation).

For mice models, 12-15 weeks old individuals (20 grams) were anesthetized through isoflurane (Forane, Baxter Health Care Corporation) inhalation (2% in oxygen for induction and 1.5% during surgery) and by subcutaneous administration of buprenorfine as analgesic. SCI surgery followed the methodology described in Reigada et al. [14]. Briefly, spinal cord was exposed by laminectomy in the 9th thoracic vertebra (T9) and subsequently received a moderate (50 KDyne) contusion using an IH Spinal Cord Impactor device.

After surgery, both rats and mice were maintained by daily manual bladder emptying for two weeks and by administration of buprenorfine as analgesic (0.03 mg/Kg), and antibiotic enrofloxazine (0.4 mg/Kg Baytril; Bayer AG) for two days.

### Neuroprotective treatments

ucf-101 (Calbiochem), an inhibitor of the pro-apoptotic protein omi/HtrA2, was administered intraperitoneally 1 h after injury and then once daily for 7 days at final concentration of 2 µmol/kg of mouse weight diluted in 100 µL saline buffer. Vehicle-treated animals received 100 µL saline buffer.

Diadenosine tetraphosphate (Ap4A; Sigma-Aldrich), a purinergic system modulator, was administered directly on the contused area of the spinal cord by placing a 1-mm^2^ piece of Spongostan (Ferrosan Medical Devices)soaked in 2 mM Ap4A. 1 h after injury and then once daily for 7 days, animals received an intraperitoneal dose of Ap4A at final concentration of 20 mg/Kg of mouse or rat weight diluted in 200 μL of saline buffer. For vehicle, animals were treated with a piece of Spongostan soaked in saline buffer and a daily intraperitoneal injection of 200 μL saline buffer for 7 days.

### Assessment of motor and sensory functions

To assess of the recovery of locomotor capacities in injured rats and mice after spinal cord injury we used the semiquantitative locomotor BBB scale (Basso, Beattie and Bresnahan locomotor scale method [16]) and BMS (Basso mice scale for locomotion method [17]), respectively. Briefly, animals were placed in an open field environment and evaluated by two observers blinded to the treatment for 4 min/animal to score locomotor abilities according to different standardized parameters.

Motor function was also evaluated according to the performance in the RotaRod test which measures the time the animal is able to remain in RotaRod 47600 device (UgoBasile) with accelerated rotational movement (from 4 to 40 rpm in 5 min test).

We also measured exercise-induced fatigue in rats using the treadmill test. The following exercise protocol was conducted in a Cinelec 21 treadmill (Cibertec): 5’ at 0.17m/s, followed by 7’30’’ at 0.25m/s, and finally 7’30’’ at 0.33m/s with an inclination of 5°. Animals were stimulated to keep moving using an electrical stimulus of 0.5 mA. Fatigue was defined as the latency time in which the animal was not accomplishing the task or was unable to keep pace with the treadmill.

Sensory function was measured using Von Frey test as described by Densmore et al. with slight modifications. Pressure nociception was measured using a Dynamic aesthesiometer 37450 to expose the animal left and right hindlimb paw to a filament executing an increasing force (from 0 to 50 grams) for 20 seconds. The time and force when the animal withdraws its paw was recorded. The same procedure was repeated in both hindlimb paws 5 times with 10 minutes resting between measurements. The mean for each paw was computed from the values after excluding the highest and lowest measurements.

### Spinal cord processing and sectioning

At the end of the experiments, 21 (mice) to 98 days (rats) after surgery, animals were euthanized with a lethal dosage of sodium pentobarbital and transcardially perfused with saline buffer with sodium heparin (1 unit/ml; Chiesi España) followed by 50-200 ml of 4% (w/v) paraformaldehyde in 0.1 M phosphate buffer pH 7.4. A segment of 1cm of spinal cord around the injured area was removed, post-fixed overnight in 4% paraformaldehyde solution (overnight at 4°C) and cryoprotected in 30% (w/v) sucrose for 1-2 days at 4°C. Finally tissue was embedded in OCT (Tissue-Tek, Sakura Finetek Europe B.V), frozen (−80°C) and in 20 µm sections using a cryostat (HM560, Microm GmbH) and mounted in microscope glass slides (Superfrost Plus Slides, ThermoFisher Scientific).

### Eriochrome Cyanine stain of myelin

To determine the area and volume of spared tissue we employed Eriochrome Cyanine staining following the protocol described by Rabchevsky and colleagues [20] modified as follows:

### Material

#### Preparation of the staining solutions

Three solutions are employed in this protocol: i) eriochrome cyanine solution, prepared by mixing 0.2 g of eriochrome cyanine RS (Sigma-Aldrich) with 0.5 mL of pure sulfuric acid (Merck) and 5 mL of 10% iron alumina solution (10 grams ferric ammonium sulfate (Sigma-Aldrich) in 100 mL of distilled water (Sigma-Aldrich)); ii) 5% iron alumina solution, a 1:1 distilled water dilution of the 10% solution; and iii) a borax solution (1 g of sodium tetraborate (Borax, Sigma-Aldrich) plus 1.25 g of potassium ferricyanide (Sigma-Aldrich) in 100 mL of distilled water. All solutions were filtered through a 3MM filter paper (Whatman) and stored in dark conditions at room temperature (RT) until use.

#### Procedure

All steps are carried out RT: i) defrost sections for 1h; ii) immerse slides in acetone for 5 min to permeabilize and fix the sections. Allow sections to air dry for 30 min; iii) introduce slides in a Coplin jar fill with filtered eriochrome cyanine solution (can be reused several times) and incubate for 30 min; iv) wash briefly by immersion in tap water (do not use distilled water in washing steps); v) immerse slides in 5% iron alumina solution for 10 (mouse) to 15 min (rat). Do not recycle the solution; vi) wash briefly by immersion in tap water; vii) immerse slides in borax solution for 5 (mouse) to 10 min (rat). Solution can be reused; viii) wash briefly by immersion in tap water; ix) proceed to dehydration by sequentially immersing the slides twice in 70% ethanol solution (diluted in distilled water, Sigma-Aldrich) for 2 min, 96% ethanol solution for 4 min, 100% ethanol solution for 8 min and HistoChoice clearing agent (Sigma-Aldrich) for 30 min; finally x) mount preparations adding a drop of DPX mounting medium (Sigma-Aldrich) on the slide and carefully place the coverslip avoiding to form air bubbles.

#### Cavalieri’s method for injury area and volume estimation

The area and volume of spared white matter were estimated from sections comprising 1cm around the epicenter (100 µm between sections), using the Cavalieri method in a Stereology Microscope (Olympus BX61) and the Olympus VIS Histoinformatics system composed by the Visiopharm Integrator System (VIS; Visiopharm) software and the NewCast module for stereology acquisition and image analysis.

Total spinal cord area and white tissue area were estimated by randomly overlaying a 2D grid of 5×5 crosses for rat and 4×4 for mouse sections on top of each section image (each cross represents 20.000 µm^2^ and 60.000 µm^2^, respectively). The number of hitting crosses was used to obtain an unbiased estimate of the white matter area. Their corresponding volumes were estimated as the multiplication of the white matter areas of the measured sections by the distance between sections, in our case 100µm. White matter area values were normalized to the values for the entire spinal cord and expressed as percentage to account for individual and positional size variability.

#### Immunofluorescence and neuronal counting

Immunofluorescence for neuronal staining and the cell counting methods were described in Reigada et al. [14]. Briefly, neurons from frozen spinal cord sections were stained by incubation with the specific neuronal marker anti-neuronal nuclei protein (NeuN, 1:500; Merk Millipore cat#MAB377, RRID: AB_11210778) diluted in blocking solution (PBS with 0.1% BSA and 0.2% Triton X100), overnight at 4°C. Then, they were incubated for 2 h at RT with an Alexa Fluor 555 nm-conjugated goat anti-mouse secondary antibody (1:500; Molecular Probes, ThermoFisher Scientific cat#A21127; RRID: AB_141596) in blocking solution. Finally, samples were stained with the fluorescent nucleic acid marker 4,6-diamino-2-fenilindol (DAPI 1:3000; Sigma-Aldrich), and mounted with a Fluorescent Mounting Medium (DAKO, Agilent Technologies Inc.). Whole spinal cord section images were captured using a Resonant Confocal microscope and analyzed using ImageJ 1.46r software (NIH; [21]) to determine the number of neuronal nuclei (see ([14] for detailed information about image processing and nuclei counting method; the employed ImageJ macro is available upon request).

#### Magnetic Resonance Imaging

Prior to sectioning and staining, (MR Acquisition type 2D) T2-weighted (T2W) axial images of fixed spinal cords from 6 male rats were acquired with the Brucker Biospec 70/30 7 Teslas MRI scanner from the experimental MRI research support facility of the Hospital Nacional de Parapléjicos. The sequence employed for the acquisition of T2-weighted images was a Bruker implementation of fast spin echo (FSE)-RARE (TurboRARE) pulse sequences. Axial spin echo images of the spinal cord were acquired with Bruker Paravision 6.0.1 software using TR/TE= 2800/22 ms; image matrix=256×256, FOV=10 mm and number of averages=5. Forty-five axial MRI slices of 0.3 mm thickness and 0.3 mm of spacing between slices, from the brain stem rostral to the C1 to the T1 vertebral level were covered by the slices.

T2 images were processed using ITK-SNAP software [22]. Briefly, hypo and/or hyperintense areas (indicating lesioned areas) were manually outlined in each slice and the outlined area measured as ITK-SNAP software. Total lesion volume was obtained by summing the areas obtained and multiplying this by the section thickness; average lesion volumes were thus obtained for each MRI paradigm. The total and the damaged volumes of each spinal cord section were exported to csv files for data analysis.

#### Data analysis

All data are expressed as mean ± SEM unless specifically stated. Statistical significance of the treatment effects was tested using unpaired Student’s t test or one-way ANOVA with Tukey post hoc test, depending to the characteristics of the data. All analyses were conducted in Prism Software 5 (GraphPad Software Inc., La Jolla, Ca, USA). Differences were considered statistically significant when p < 0.05.

## RESULTS AND DISCUSSION

To evaluate the accuracy of the Cavalieri’s estimator in the assessment of cellular and pathophysiological outputs induced by spinal cord injury (SCI) we used previously published and ongoing experimental data to estimate the potency and limitations of stereological methods as biomarker for the diagnostic and prognostic for experimental SCI. We analysed the correlation of stereology data obtained from eriochrome cyanide stained sections with anatomical, motor and sensorial data in rat and mouse models of contusion.

### The Cavalieri’s estimator and the measurement of neurodegeneration

EC-Cavalieri’s method is designed to estimate total and white matter areas and volumes present in each section or region of spinal cord, respectively. Reduction in white matter areas and volumes are employed to estimate injury extension. It is, thus, an indirect measurement of injury size. Magnetic resonance imaging (RMI) is an alternative method that directly measures the injury size which is routinely employed in the clinical practice as a diagnostic and prognostic marker of spinal cord injury. Previous analyses have demonstrated a good agreement between MRI and histological estimates [23]. To evaluate the agreement specifically with Cavalieri’s estimations, we have compared *ex vivo* MRI’ and Cavalieri’s estimates of total tissue and tissue damage –areas and volumes– in 5 fixed spinal cords sampled 8 weeks after a mild contusive SCI (150Kdynes) in T8 vertebra. The section or slice of minimal tissue preservation (total tissue for MRI and WM for Cavalieri) was identified as the injury epicenter and employed to determine the relative positions of the remaining sections and slices.

The comparisons of total area estimated after MRI and Cavalieri methods reveal a good agreement in the profiles of each spinal cord (Fig 1A).

**Figure 1:**
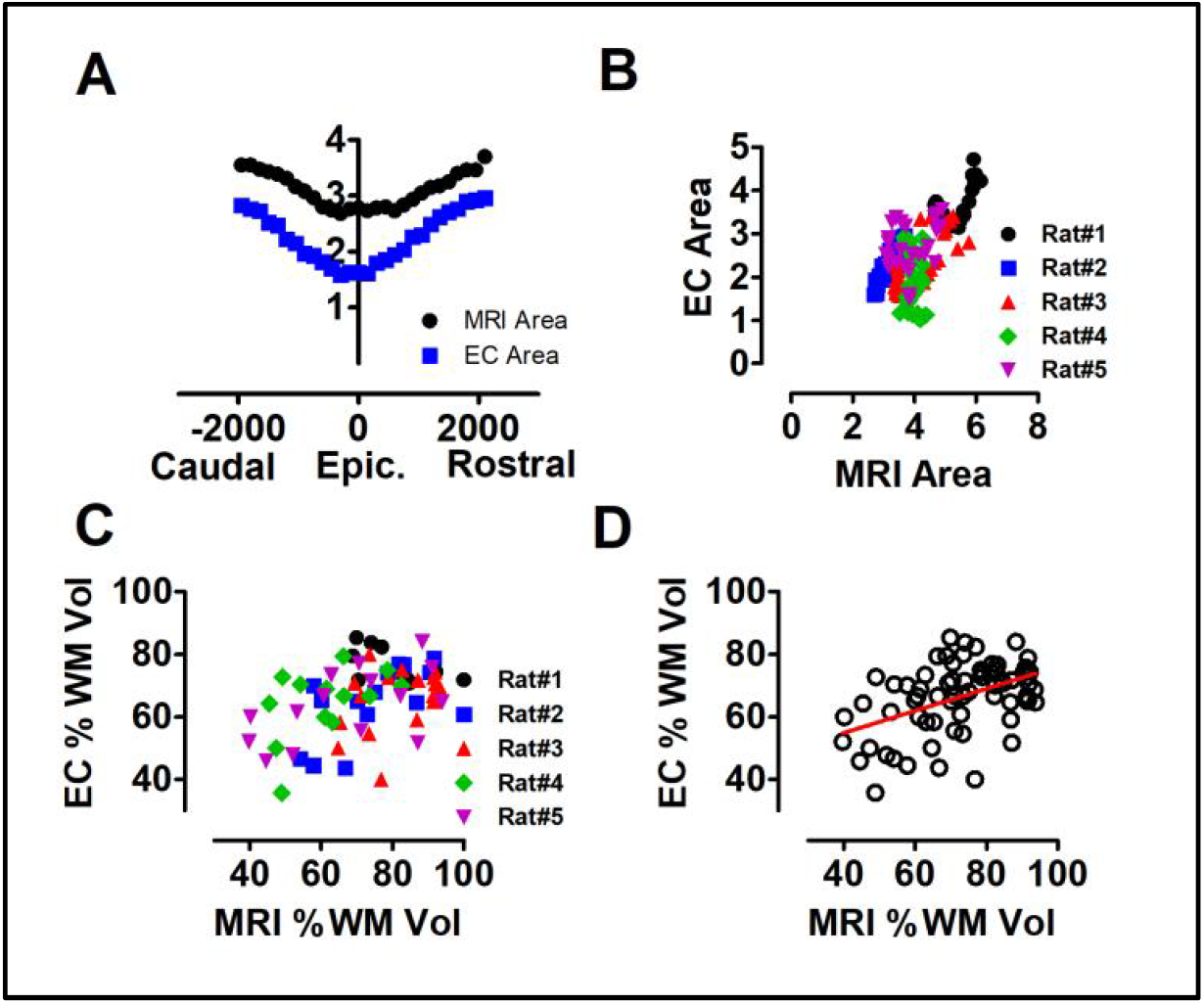
Cavalieri’s estimations highly correlate with MRI-estimated neurodegeneration and neuronal loss. **A**. Comparison of the axial area estimations obtained from MRI and Cavalieri’s in individual #2. The dot plot represents in the X-axis the distance to the epicenter (in microns) and in the y-axis the individual area of each section or slice. Area values correspond to the simple moving average of 5 values to smooth the curves and interpolate non available measures. **B**. Correlation between MRI and Cavalieri’s estimates of axial area (R=0.6834, p<0.001). Analyzed individuals are indicated. Data correspond to simple moving average of 5 raw values. **C**. Comparison of the volume of preserved tissue obtained from MRI (%preserved tissue) and the Cavalieri’s (% of preserved WM) across the 4 individuals. Anomalous individual #4 (green) was excluded from Pearson correlation (Pearson R=-0.825, p=0.175, n=4). **D**. Correlation between MRI and Cavalieri’s estimates of preserved tissue in the corresponding slices and section (R=0.4674, p<0.001, n=70). Analyzed individuals are indicated.

Agreement is confirmed by Pearson’s correlation across the different spinal cords (R=0.6834, p<0.001, Fig 1B). Despite the overall similarity, one individual (spinal cord 4) shows large discrepancy between estimates, with the areas estimated from MRI remaining fairly constant along the spinal cord whereas histology indicates a clear reduction in the sections surrounding the epicenter. Other obvious differences involve the size and variability of the estimated areas, with MRI estimations providing larger and less variable values than Cavalieri’s estimations. Size differences likely result from the shrinking of the tissue during histological processing whereas the larger variability in Cavalieri’s estimates is a logical consequence of the damages and folds commonly occurring during histological processing.

Concerning the quantification of the damaged tissue, identifying its precise contour on the MRI images was difficult, sometimes arbitrary, due to absence of information that clearly delimitates the spared from the damaged tissue. On the contrary, EC staining clearly differentiates the spared white matter from the rest of the tissue. Interestingly, despite these difficulties and the presence of the anomalous individual 4, a fairly good agreement can be observed between the injury volume estimated by MRI and the preservation of white matter estimated using Cavalieri (Pearson R=-0.825, p=0.175, n=4; Fig 1C). When analyzed in individual sections/slices, values show a lower positive correlation (R=0.4674, p<0.001, n=70; Fig. 1D) though highly significant due to the increased number of comparison available.

As a whole, comparison indicates that Cavalieri’s estimations broadly agree with MRI measurements. However, these are preliminary analyses that can be much improved by increasing sample size and using more specific tools; however, they illustrate the capability of Cavalieri’s method on EC stained sections to estimate injury size. Similar agreement between histological and MRI estimations has been previously described [23].

As we have already mentioned, Cavalieri’s method provides an unbiased estimate of white matter preservation that can be employed to estimate injury size but, does it reflect gray matter loss, or more specifically, the loss of gray matter neurons?. To answer this question, we studied the correlation between the extension of the damaged tissue estimated using Cavalieri and number of surviving neurons using data published in Reigada et al. [14].

In this article, both injury volume and number of preserved neurons were measured in sections from mice spinal cords sampled 21 days after a moderate SCI contusion (50 Kdynes). The analysis of this data shows that the higher tissue damage the higher neuronal loss (Figs. 2A and 2B) and, not significantly, both parameters have a high degree of correlation (Pearson’s R = -0.57; *p*=0.12, n=6). It also highlights a major difference in the estimated extent of neurodegeneration reflecting the massive neuronal loss that extends throughout the analysed range of distances (1.2 mm caudal and rostral for the injury epicenter) characteristic of the subacute phase of SCI while white matter degeneration is less extensive (spared WM remains in every analysed section and normal values are observed 1 mm away the epicenter) and proceeds more gradually [4].

**Figure 2:**
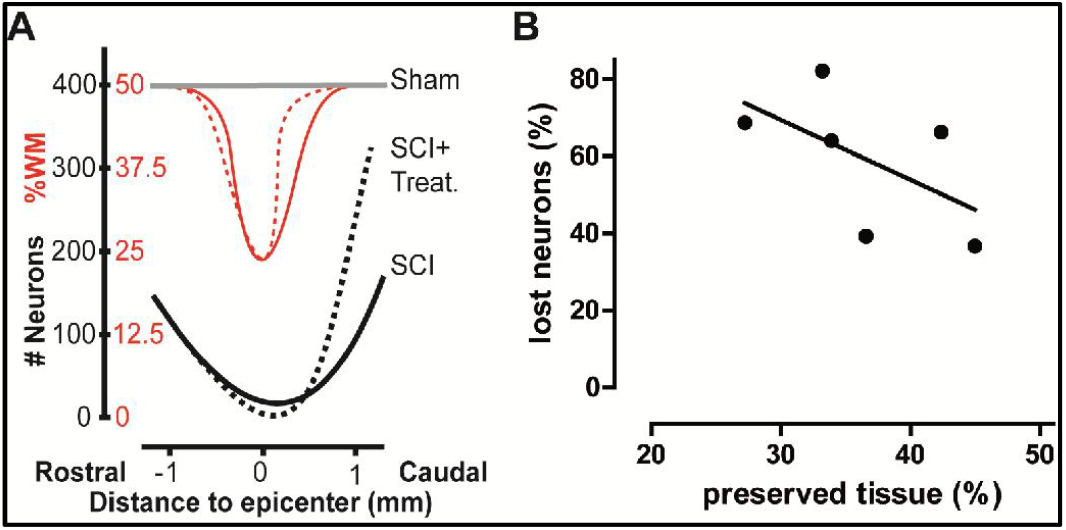
Estimation of tissue damage degree highly correlates with neuronal cell death levels after moderate contusion SCI. **A**. Spatial changes in spared white matter and neuronal abundance in undamaged and contused (treated and untreated) spinal cords. Adapted from Reigada *et al* [14]. **B**. Correlation between volume of spared white matter estimated by the Cavalieri stereological method and percentage of lost neurons. Each data point corresponds to one individual mouse after 21 days of a moderate contusion in T9 spinal vertebra. Line represents the linear regression fit of the data and shows a high degree correlation, where the increase in tissue damage (lower % of white matter (WM)) is related to an increase of the percentage of neuronal cell loss (Pearson’s coefficient = -0.57; *p*=0.12, n=6).

### Cavalieri’s estimations predict sensory and motor recovery

In SCI, the extent of neurodegeneration determines the resulting functional deficits. Consequently, EC-Cavalieri’s estimations should be expected to inform on the extent of the functional loses caused by the SCI. Motor function in murine models of SCI is measured through different test, among which the BBB and BMS scales for rats and mice, respectively, are the most broadly employed ([16, 17]). In these tests, researchers score multiple parameters of the animal’s locomotion in an open field, including joint movement of the hind limbs, weight support, stepping, coordination, trunk stability, and tail position. Afterwards, the so-obtained values are employed to compute a score and a subscore.

Comparison of Cavalieri’s estimates with BMS scores in mice 7 days after a moderate contusion (50 KDynes) reveals a significant correlation (Fig. 3A_i), with higher scores in individuals with smaller injuries (Pearson’s R coefficient=0.56; *p*=0.045*, n=10). However, 21 days after injury (days post-injury, dpi) the correlation becomes lost because all individuals present similar BMS scores irrespectively of their WM preservation (Fig. 3A_iii). When we compare Cavalieri’s estimations with the BMS subscores–a different scoring approach of the same parameters–the reverse is observed, with a higher correlation at 21 dpi (Fig. 3A_iv; Pearson’s R coefficient=0.38; *p*=0.14, n=10) than at 7 dpi when almost all individuals have subscores equal to 0 (Fig. 3A_ii).

**Figure 3:**
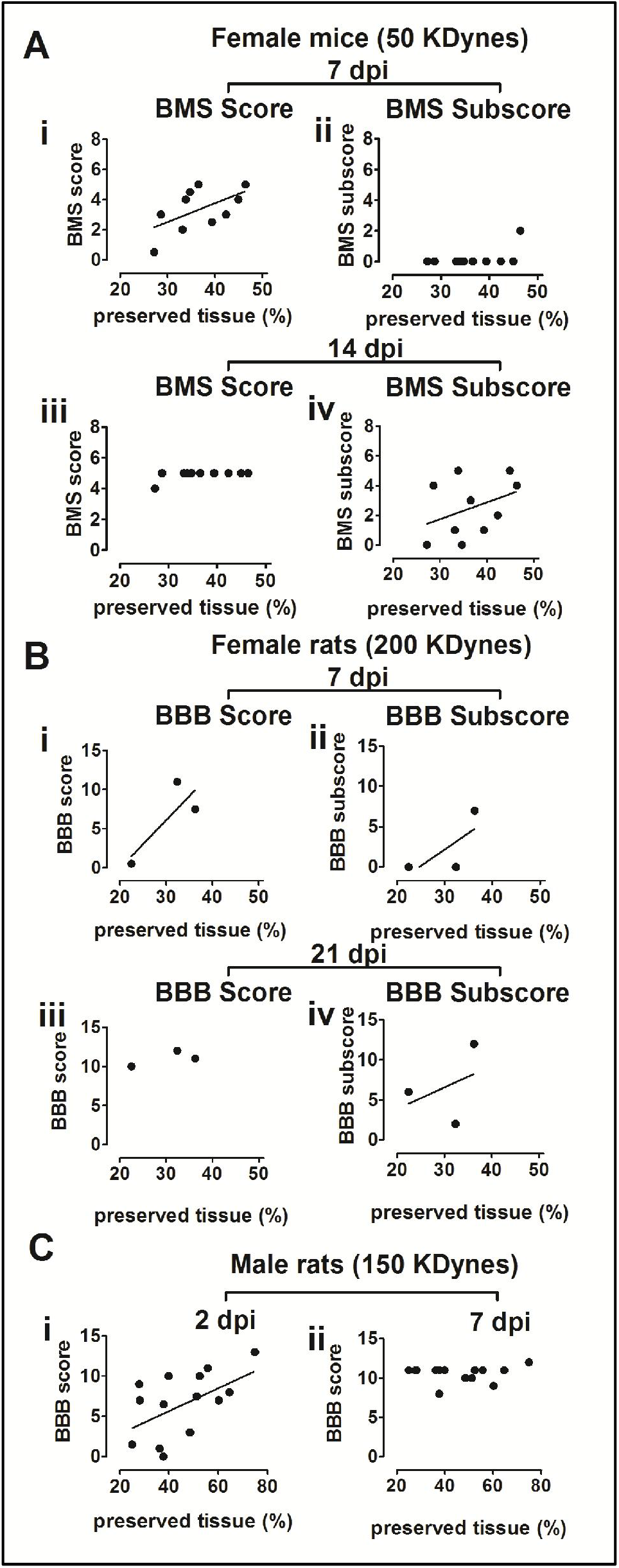
Estimation of tissue damage degree correlates with locomotor scale data at early times after injury in both mouse and rat SCI models. Dot plot summary of the correlation between volume of spared white matter estimated by the Cavalieri stereological method and the different scores that estimates the locomotor recovery capacity after SCI. Each data point corresponds to one individual and line represents the linear regression fit of the data. Data analysis shows that tissue damage decrease correlates with a global locomotor recovery capacity at early stages of SCI: 7 dpi in moderate SCI in mice (**A_i**; Pearson’s coefficient= 0.56; *p*=0.045*, n=10) and 2 dpi in mild SCI in rats (**C_i**;Pearson’s coefficient at 2 dpi = 0.52; *p*=0.02*; n=14). Low number of rats with a moderate SCI analysed (**B_i**) does not allow to statistically fit the data but the tendency is the same than in mice. These correlations are lost at later times: 21 dpi in mouse moderate SCI (**A_iii** and **B_iii**) and 7 dpi in rat mild SCI (**C_ii**). Subscores are a more detailed analysis and estimates the onset of keystone capacities for regular locomotion (subscores). These appeared at later times after 21 days of a moderate SCI and also correlate with tissue damage degree in both mice (**A_iv**; Pearson’s coefficient at 21 dpi = 0.38; *p*=0.14, n=10) and rats (**B_ii and B_iv**).

Similar results are observed when analyzing rats following moderate injury (200 KDynes). Due to the very small population available for this study (n=3), results are not conclusive, but there is a clear correlation between BBB score and WM preservation during the first days (Fig. 3B_i) that is lost at 21 dpi when the 3 individuals show similar BBB scores (Fig. 3B_iii). BBB subscores also perform similar to mice, with a better correlation at 21 dpi (Fig. 3B_ii) than at 7 dpi (Fig. 3B_iv).

When analysing a milder injury (150 Kdynes) in rats, the resulting pattern is similar, both the highest correlation and its decline are observed earlier –2 dpi for maximum correlation(Fig. 3C_i; Pearson’s coefficient = 0.52, *p*=0.02*, n=14), and loss of correlation at 7 dpi (Fig. 3C_ii)–. These repeated temporal patterns in the correlation between WM preservation and motor function primarily reflects the nature of the BBB and BMS scales which mainly score the achievement of certain functional keystones. In the experiments here presented, both mice and rats rarely achieve hind-forelimb coordination (BBB score=11, BMS score=5) until much later than the times analysed causing the animals to reach the same scores despite their different functional deficits (and WM preservation).

Alternative locomotor evaluations –like the RotaRod or Treadmill tests– provide greater sensitivity and more linear data than BBB and BMS scales, although they can not be used in non-walking animals. RotaRod test evaluates motor coordination and balance, two motor skills that are the sum of a set of functions owned in an specific moment. When compared to Cavalieri’s estimates, RotaRod values from mice with moderate contusion (50 KDynes) show a high correlation that is sustained at the studied time(Fig. 4A; 14 dpi: Pearson’s R coefficient=0.62; *p*=0.07; 21 dpi: Pearson’s coefficient = 0.49; *p*=0.13; n=7). Treadmill test –a measurement of the time the animal can sustain locomotion under defined conditions of speed and inclination as a measurement of fatigue– also shows a good correlation with WM preservation in rat 28 days after a mild (150 Kdynes) contusion(Fig. 4B; Pearson’s coefficient = -0.4, *p*=0.07; n=15).

**Figure 4:**
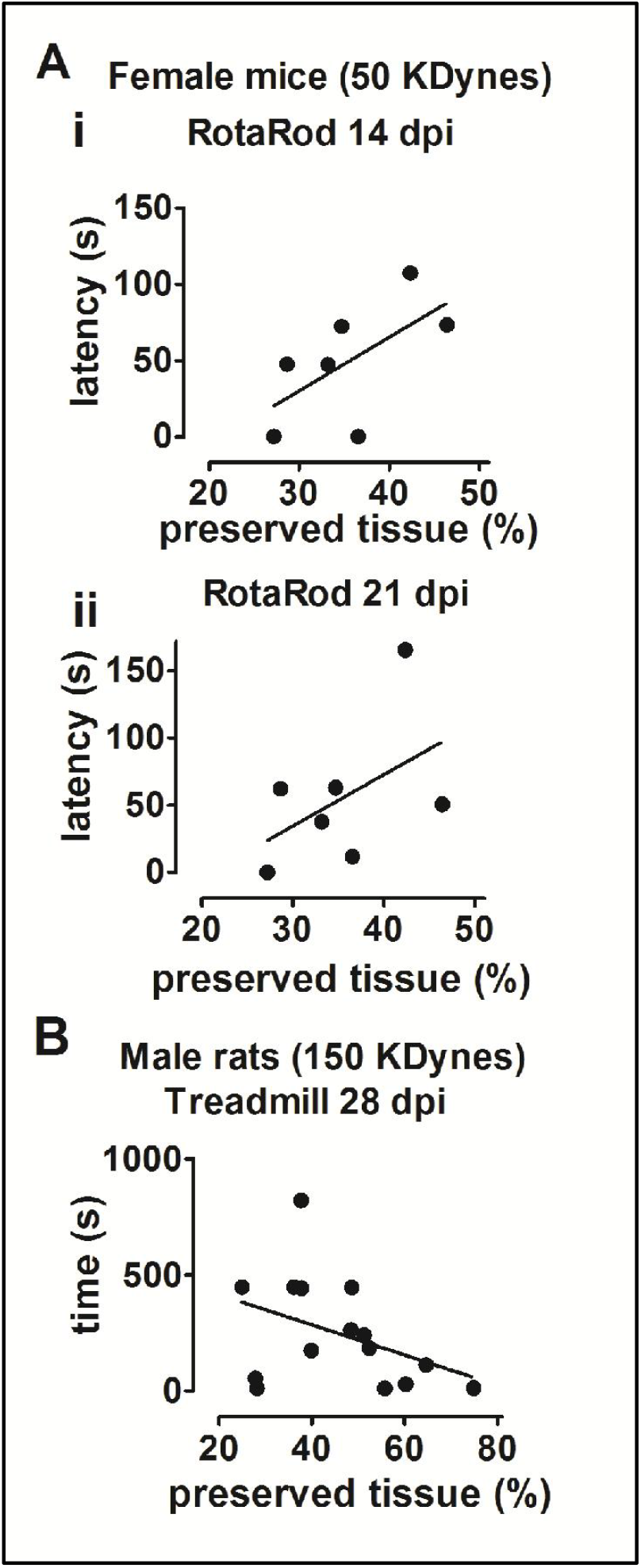
Estimation of tissue damage degree highly correlates with motor capacities data in both mouse and rat SCI models. Dot plot summary of the correlation between volume of spared white matter estimated by the Cavalieri stereological method and the locomotor performance after SCI estimated by the RotaRod and Treadmill methods. Each data point corresponds to one individual and line represents the linear regression fit of the data. In mice, after moderate SCI (**A**), there is a high degree of correlation where tissue damage decreases the global locomotor skills measured by latency time in RotaRod at14 dpi (Pearson’s coefficient = 0.62; *p*=0.07, n=7) and21 dpi Pearson’s coefficient = 0.49; *p*=0.13, n=7). In rats after mild SCI (**B**) there is also a high degree of correlation where tissue damage decreases the global locomotor skills measured by Treadmill at 28 dpi (Pearson’s coefficient = -0.4; *p*=0.07, n=15)

We also tested how Cavalieri’s estimations relate to sensory deficits using the Von Frey method. This method is a mechanical nociceptive threshold test that provides a range of forces to the hind paws in order to find the force at which the subject reacts because the sensation is painful.

As observed for motor function, von Frey values shows a significantly high correlation with tissue damage in the same set of rats mildly (150 KDynes) contused at 28 dpi (Fig. 5), with a decrease in pain sensation as volume of tissue damage increases (Pearson’s R coefficient =-0.68, *p*=0.029; n=15).

**Figure 5:**
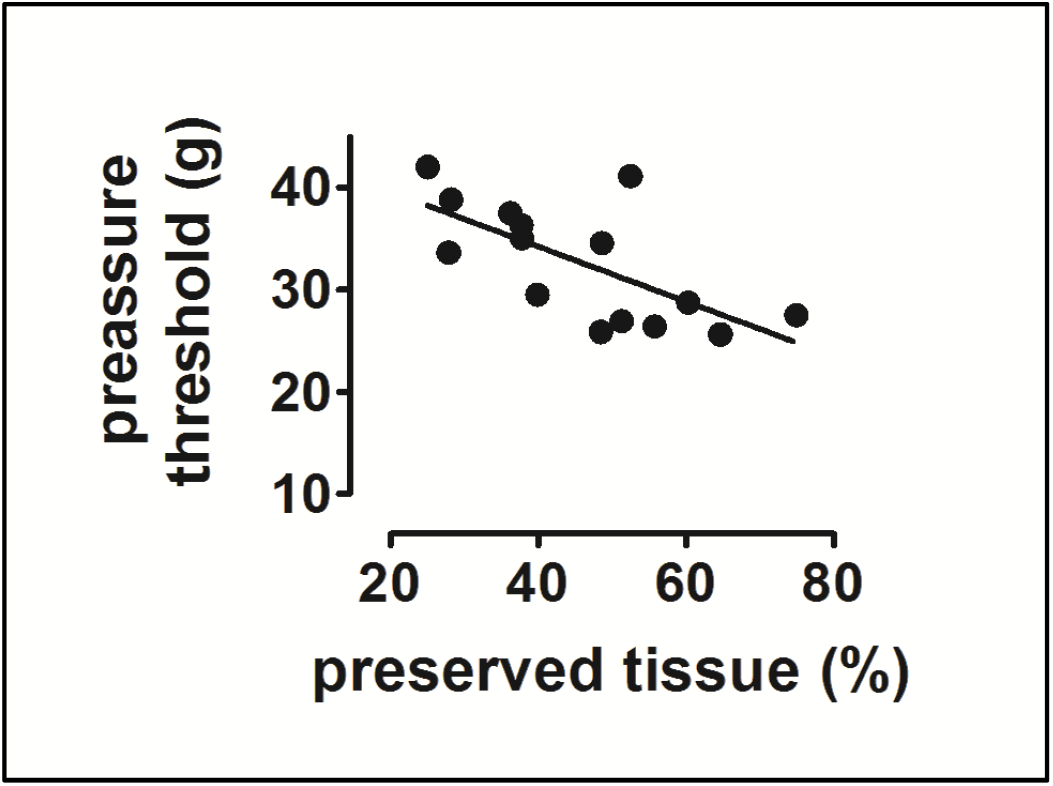
Estimation of tissue damage degree highly correlates with sensorial capacities data in rat SCI model. Dot plot summary of the correlation between volume of spared white matter estimated by the Cavalieri stereological method and the sensorial threshold evaluated through by the von Frey method. Each data point corresponds to one individual and line represents the linear regression fit of the data. After mild SCI in male rats, there is a high degree of correlation, a tissue damage decrease and an increase in sensitivity at 28 dpi (correlation: Pearson’s coefficient = -0.68; *p*=0.029, n=15).

### Cavalieri’s estimates inform on the effects of neuroprotective treatments

To explore the potency of Cavalieri’s estimations in the study of the effects of neuroprotective treatments efficacy, we employ the data from two published studies ([13,14]) in which we tested neuroprotective therapies to reduce apoptotic (ucf-101) and excitotoxic (Ap4A) neuronal death in a mouse model of SCI by moderate contusion (50 KDynes).

In both studies, the treatments did not reduce significantly the total volume of damage measured 21 dpi (Fig. 6A and 6B, see also Fig 2B) but the analysis of individual sections revealed that both treatments did significantly reduce tissue loss in caudal regions. Therefore, Cavalieri’s estimations of tissue damage indicate that both treatments protect neural tissue in regions caudal to the injury epicenter. These results are confirmed by the analyses of neuronal abundance across the spinal cord after treatment with ucf-101. As illustrated in Fig. 6C_i, the whole damaged spinal cord region shows a big dispersion in neuronal counts and no differences in the total number of surviving neurons are found between vehicle and ucf-101 treated animals. However, when we analysed only caudal sections (Fig. 6C_ii), the observed increase in tissue preservation is highly related to increased neuronal survival. The impact of tissue preservation in caudal sections is appreciated in the locomotor function recovery. When we analyse the effects of ucf-101 and Ap4A neuroprotective treatments on the previously described relation between Cavalieri’s estimations and BMS scores (see Fig. 3), we can observe changes in that relationship (Fig. 7A_i and 7B_i) resulting from the overcoming of the locomotor limits (vg in the hindlimb forelimb coordination) observed in control animals at 21 dpi.

**Figure 6:**
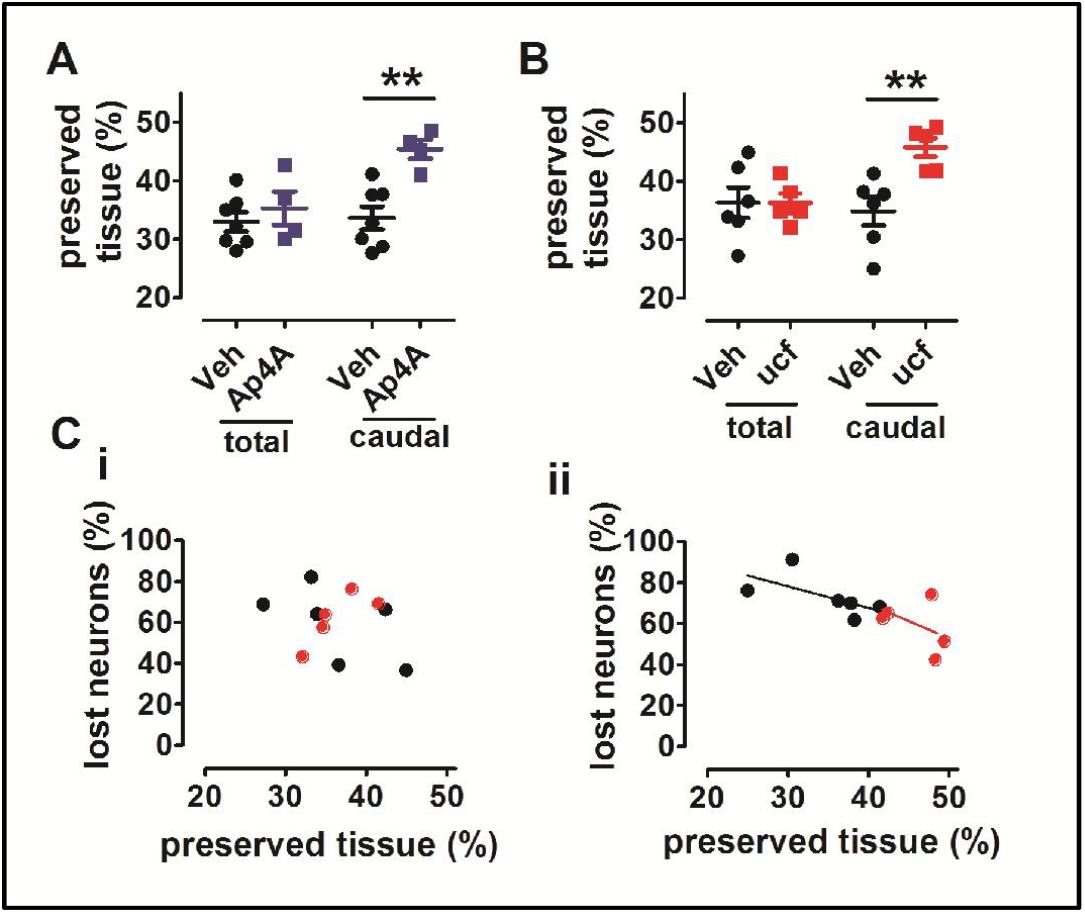
Neuroprotective treatments induce a reduction in tissue damage limited to caudal areas of the injury, and correlates with a decrease in neuronal cell death. Dot plot summary of the correlation between volume of spared white matter estimated by the Cavalieri’s stereological method and the neuronal cell death percentage. Each data point corresponds to one individual mouse after 21 days of a moderate contusion in T9 spinal vertebra. Black symbols represent vehicle-treated animals and colour symbols ucf-101 (red) and Ap4A (blue) treated animals. Lines represents the linear regression fit of the vehicle (black) and drug (red) treated animals. Both treatments reduce tissue damage but limited to caudal sections of the injury (**A** and **B**) and it correlates with the reduction of neuronal death also limited only to caudal sections(total volume **C_i** vs caudal volume **C_ii**). (Pearson’s coefficient = -0.43; *p*=0.23, n=5).

**Figure 7:**
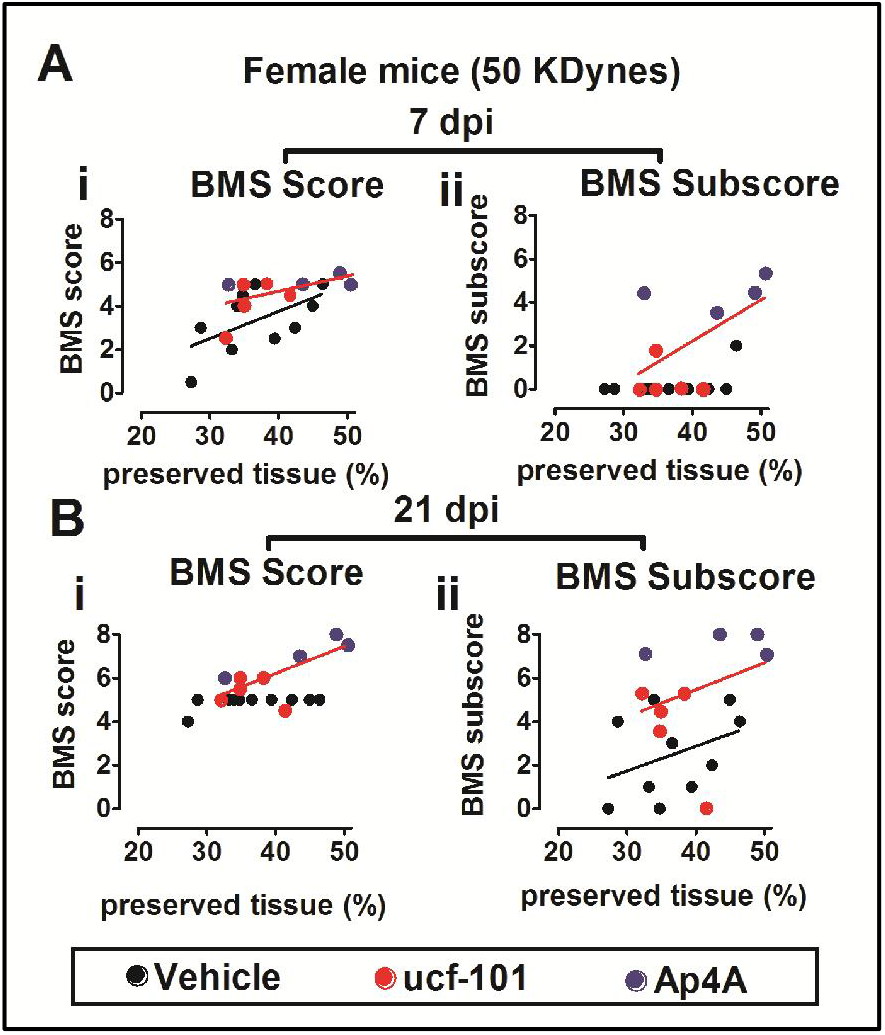
Neuroprotective treatments in mouse SCI model induce an improvement of locomotion, reducing time onset of keystone parameters of walk and increasing duration and degree of the recovery. Dot plot summary of the correlation between volume of spared white matter estimated by the Cavalieri’s stereological method and the different scores that estimates the locomotor recovery capacity after SCI. Each data point corresponds to one individual mouse after 21 days of a moderate contusion in T9 spinal vertebra. Black symbols represent vehicle-treated animals and colour symbols ucf-101 (red) and Ap4A (blue) treated animals. Lines represent the linear regression fit of the vehicle (black) and drug (red) treated animals. Correlation analysis of score data shows that treatments induce an elongation of the recovery time, further the 7 dpi limit of vehicle-treated animals (**A_i**), reaching scores above the stabilization level showed by vehicle-treated animals 21 dpi (**B_i**) and strongly correlates with the estimated tissue damage.(at 7 dpi: Pearson’s coefficient = 0.56; *p*=0.06, n=9 / at 21 dpi: Pearson’s coefficient = 0.72; *p*=0.013, n=9;). In the case of subscore, important keystone parameters of locomotion are reached earlier than in vehicle-treated animals (7 dpi; **A_ii**) and are maintained higher a longer time until 21 dpi (**B_ii**) with a high correlation with stereological data (at 7 dpi: Pearson’s coefficient = 0.58; *p*=0.05, n=9; / at 21 dpi: Pearson’s coefficient = 0.32; *p*=0.2, n=9).(Vehicle data is previously shown in Fig.2)

The same applies to BMS subscore data, in which neuroprotection allows locomotor improvements at 7dpi (Fig. 7A_ii) that are not present in control animals (Fig.3) until 21 dpi (Fig.7B_ii) leading to a sustained correlation (7 dpi:BMS score Pearson’s R coefficient=0.56; *p*=0.06, n=9; BMS subscore: Pearson’s R coefficient=0.58; *p*=0.05, n=9; at 21 dpi: BMS score: Pearson’s R coefficient=0.72; *p*=0.013, n=9; BMS subscore: Pearson’s R coefficient=0.32; *p*=0.2, n=9; Control data are shown in figure 3).

## Conclusions

The review of the present evidence confirms EC Cavalieri’s estimations of white matter loss as a surrogate of neurodegeneration in SCI both at cellular and tissue levels. EC-Cavalieri’s estimations also predict functional recovery and provide key information on the responses to treatments.

## Acknowledgements

We thank the Fundación del Hospital Nacional de Parapléjicos para la Investigación y la Integración (FUHNPAIIN) and the MRI, microscopy and animal facilities from the Research Unit of the Hospital Nacional de Parapléjicos (Toledo, Spain) for their technical and logistic support. This work was funded by: Fundación para la Investigación Sanitaria de Castilla la Mancha (FISCAM; PI-2010/19), Consejería de Sanidad de Castilla la Mancha (PI-06066-00), Fondo de Investigación Sanitaria, Instituto Carlos III (FIS-ISCIII; PI-081941), International Foundation for Research in Paraplejía (IRP; P181), SAMOS Medical Enterprise SL, and co-financed by the European Union (FEDER) “A way to make Europe”.

## Notes

### Competing Interest Statement

The authors have declared no competing interest.

